# End-to-end multitask learning, from protein language to protein features without alignments

**DOI:** 10.1101/864405

**Authors:** Ahmed Elnaggar, Michael Heinzinger, Christian Dallago, Burkhard Rost

**Affiliations:** TUM (Technical University of Munich) Department of Informatics, Bioinformatics & Computational Biology - i12, Boltzmannstr. 3, 85748 Garching/Munich, Germany; TUM Graduate School, Center of Doctoral Studies in Informatics and its Applications (CeDoSIA), Boltzmannstr. 11, 85748 Garching, Germany; Institute for Advanced Study (TUM-IAS), Lichtenbergstr. 2a, 85748 Garching/Munich, Germany & TUM School of Life Sciences Weihenstephan (WZW), Alte Akademie 8, Freising, Germany & Columbia University, Department of Biochemistry and Molecular Biophysics, 701 West, 168^th^ Street, New York, NY 10032, USA

**Keywords:** Machine Learning, Language Modelling, Semi-Supervised Learning, Multi-Task Learning, Protein Secondary Structure Prediction, Protein subcellular-localization Prediction

## Abstract

Correctly predicting features of protein structure and function from amino acid sequence alone remains a supreme challenge for computational biology. For almost three decades, state-of-the-art approaches combined machine learning and evolutionary information from multiple sequence alignments. Exponentially growing sequence databases make it infeasible to gather evolutionary information for entire microbiomes or meta-proteomics. On top, for many important proteins (e.g. *dark proteome* and *intrinsically disordered proteins*) evolutionary information remains limited. Here, we introduced a novel approach combining recent advances of Language Models (LMs) with multi-task learning to successfully predict aspects of protein structure (secondary structure) and function (cellular component or subcellular localization) without using any evolutionary information from alignments. Our approach fused self-supervised pre-training LMs on an unlabeled big dataset (UniRef50, corresponding to 9.6 billion words) with supervised training on labelled high-quality data in one single end-to-end network. We provided a proof-of-principle for the novel concept through the semi-successful per-residue prediction of protein secondary structure and through per-protein predictions of localization (Q10=69%) and the distinction between integral membrane and water-soluble proteins (Q2=89%). Although these results did not reach the levels obtained by the best available methods using evolutionary information from alignments, these less accurate multi-task predictions have the advantage of speed: they are 300-3000 times faster (where HHblits needs 30-300 seconds on average, our method needed 0.045 seconds). These new results push the boundaries of predictability towards grayer and darker areas of the protein space, allowing to make reliable predictions for proteins which were not accessible by previous methods. On top, our method remains scalable as it removes the necessity to search sequence databases for evolutionary related proteins.

## Introduction

### Successful combination of evolutionary information and artificial intelligence

Predicting structural and functional aspects of proteins based on their amino acid sequence has been one of the most challenging problems for computational biology. Researchers have been working on this problem for nearly five decades in order to bridge the sequence-structure/function gap, i.e. the gap between 180M proteins of known sequence (UniProt (Consortium, 2018)) and about 560K proteins with experimental annotations about function (SwissProt (Boutet, et al., 2016)) or 150K with high-resolution experimental structures (PDB (Berman, et al., 2000)). The biggest single improvement in prediction performance was achieved over two decades ago through the combination of machine learning (ML) and evolutionary information, i.e. the profiles extracted from <u>Multiple Sequence Alignments (MSA)</u> of related proteins as input feature for protein secondary structure prediction (Rost and Sander, 1993; Rost and Sander, 1993; Rost and Sander, 1994). Evolutionary information was quickly adopted by the field (Frishman and Argos, 1995; Jones, 1999; Mehta, et al., 1995) and has become the *de facto* standard for encoding protein sequences for most machine learning applications, including the prediction of transmembrane helices (Rost, et al., 1995), solvent accessibility (Pollastri, et al., 2002; Rost and Sander, 1994), protein flexibility (Capriotti, et al., 2005; Radivojac, et al., 2004), inter-residue contacts (Hayat, et al., 2015), protein-protein interactions (Hamp and Rost, 2015; Zhang, et al., 2012), and subcellular localization (Casadio, et al., 2008; Goldberg, et al., 2014; Nair and Rost, 2003).

With protein sequence data exploding exponentially, evolutionary information becomes even more valuable (Jones, 1999; Rost, 2001). However, even today’s fastest solutions, HHBlits3 (Steinegger, et al., 2019) and MMSeqs2 (Steinegger and Söding, 2017), can hardly cope with all 2.5B sequences in BFD (Steinegger, et al., 2019). On top, evolutionary information is more tricky to obtain for proteins that are difficult to align such as intrinsically disordered proteins (IDPs), or proteins from the Dark Proteome (Perdigao, et al., 2015).

### Mining the wealth of unlabeled bio-data through transfer learning

What if we could replace the search for evolutionary related proteins by semi-supervised multi-task (MT) learning? Such an approach would build upon the recent success of language models (LMs) (Devlin, et al., 2018; Howard and Ruder, 2018; Peters, et al., 2018) applied to protein sequences (Heinzinger, et al., 2019; Rives, et al., 2019). Language models are trained on large unlabeled text-corpora to predict the most probable next word in a sentence, given all previous words in this sentence (auto-regression). As only a sequence of words (sentence) is needed to train, such approaches are referred to as *self-supervision*, i.e. the learning of syntax and semantics through data without requiring expert knowledge. The same idea can be adopted to computational biology by considering single amino acids as words and their sequences as sentences. This simple transfer unleashes the power of big data sets easily outgrowing text-corpora used in NLP such as Wikipedia by orders of magnitude (Heinzinger, et al., 2019; Howard and Ruder, 2018; Rives, et al., 2019).

So far applications of LMs to protein sequences have split the training into two phases (Alley, et al., 2019; Heinzinger, et al., 2019; Howard and Ruder, 2018): (1) self-supervised pre-training on unlabeled data learns vector representations (embeddings), which is followed by (2) supervised training of a second network on labeled data using the embeddings as input. Although transfer learning succeeds to an amazing extent to extract some principles relevant for the understanding the language of life, databases of protein sequences do not capture any explicit information about the molecular constraints shaping proteins in evolution. Not surprisingly, the simple two-step solution self-supervised transfer-learning followed by supervised deep-learning fails to beat the best methods using evolutionary information. For NLP, multi-task (MT) learning merges the two steps in an end-to-end trainable machine, e.g. through transformer models currently reaching the state-of-the-art in NLP (Dai, et al., 2019; Devlin, et al., 2018; Radford, et al., 2019).

Here we hypothesized that multi-task (MT) learning into one end-to-end network might enable to fine-tune NLP models to particular supervised problems in computational biology. More specifically, we trained an MT transformer model on five different tasks: (1) The language modeling task was trained on 35M sequences from UniRef50 (Suzek, et al., 2015); (2) The per-residue (word-level) task was the prediction of secondary structure in three (and eight) states derived from DSSP (Kabsch and Sander, 1983); (3) The per-protein (sentence-level) tasks included the predictions of protein subcellular localization in ten classes and a binary classification into membrane-bound and soluble proteins. In order to simplify the comparability of results between different approaches, we used two datasets from recent state-of-the-art publications (Almagro Armenteros, et al., 2017; Klausen, et al., 2019).

## Materials & Methods

### Data

#### Language modeling

The self-supervised, language modelling task was trained on UniRef50 a subset of UniProt (Consortium, 2018) clustered at 50% pairwise sequence identity (PIDE). The database contained 33M protein sequences with 9,577,889,953 residues and a vocabulary of 25 words: the standard 20 and two rare amino acids (U and O) along with three letters for ambiguous (B, Z) or unknown amino acids (X). Each protein was treated as a sentence and each amino acid was interpreted as a single word. The model was trained using 99.9% of the data, while randomly keeping 0.01% (∼33k) of the proteins for validation. In this case, since the goal is self-supervision, homology doesn’t play a major role and thus train/validation splits could be drawn at random. The maximal model length was set to 1024; for proteins longer than 1024 residues only the first 1024 residues were used.

#### Per-residue: secondary structure

Using a data set of protein secondary structure previously published (Klausen, et al., 2019) simplified comparisons of this task to state-of-the-art prediction methods, including: NetSurfP2.0 (Klausen, et al., 2019), Spider3 (Heffernan, et al., 2017), RaptorX (Wang, et al., 2016) and JPred4 (Drozdetskiy, et al., 2015). For training, 10,837 sequence-unique (at 25% PIDE) proteins with high-quality 3D structures (<2.5Å or 0.25nm) were collected by the PISCES server (Wang and Dunbrack Jr, 2003) from the Protein Data Bank – PDB (Berman, et al., 2000). DSSP (Kabsch and Sander, 1983) assigned secondary structure; residues without atomic coordinates (*REMARK-465*) were removed. The seven DSSP states (+ one for unknown) were mapped to three states as in previous works (Rost and Sander, 1993): [G,H,I] → H: helix, [B,E] → E: strand, all others to O: other). Since NetSurfP-2.0 only published pre-computed profiles, protein sequences were extracted through SIFTS (Velankar, et al., 2012). During this mapping step, 56 proteins were removed from the training and three from the test set due to differences in lengths between SIFTS and NetSurfP-2.0 (two from CB513 (Cuff and Barton, 1999); one from CASP12 (Abriata, et al., 2018); none from TS115 (Yang, et al., 2016)). Proteins longer than 512 residues were also removed due to constraints of our transformer models. Three test sets (also referred to as validation sets) were distinguished: TS115: 115 proteins from high-quality structures (<3Å) released after 2015 (and at most 30% PIDE to any protein of known structure in the PDB at the time); CB513: 513 non-redundant sequences compiled 20 years ago (511 after SIFTS mapping); CASP12: 21 proteins taken from the CASP12 free-modelling targets (20 after SIFTS mapping; all 21 fulfilled a stricter criterion toward non-redundancy than the two other sets; non-redundant with respect to all 3D structures known until May 2018 and all their relatives). The NetSurfP-2.0 authors removed all proteins from training with >25% PIDE to any protein in the test sets.

#### Per-protein data: subcellular localization and membrane/soluble

The prediction of subcellular localization (also referred to as *location* or *cellular component*) was trained and evaluated through data from the authors of DeepLoc (Almagro Armenteros, et al., 2017), one of the state-of-the-art prediction methods for this task. The performance of several other methods was also evaluated on this set, namely: LocTree2 (Goldberg, et al., 2012), MultiLoc2 (Blum, et al., 2009), SherLoc2 (Briesemeister, et al., 2009), CELLO (Yu, et al., 2006), iLoc-Euk (Chou, et al., 2011), WoLF PSORT (Horton, et al., 2007) and YLoc (Briesemeister, et al., 2010). This data set pools proteins with experimental annotation (code: ECO:0000269) from UniProtKB/Swiss-Prot (release: 2016_04, (Consortium, 2018)). To facilitate training and evaluation, the authors of DeepLoc mapped the plain localization annotations to ten classes, removing all proteins with multiple annotations. Additionally, all proteins were labelled as *water-soluble* or *membrane-bound* (some with ambiguous annotations as *unknown*). The resulting 13,858 proteins clustered into 8,464 representatives with PSI-CD-HIT (Fu, et al., 2012; Li and Godzik, 2006) (v4.0; at 30% PIDE or E-val<10-6; for alignments covering>80% of the shorter protein). The remaining proteins were split into training and testing by using the same proteins for testing as DeepLoc. Similar to DeepLoc, proteins longer than the maximum processed by the method (1000 for DeepLoc, 1024 for the method presented here) remained in the data set and were simply cut (keeping the first and the last 512 residues).

#### Evaluation measures

For simplicity of comparison, the evaluation measures followed previous publications (Almagro Armenteros, et al., 2017; Klausen, et al., 2019), namely three-state per-residue accuracy (<u>Q3</u> (Rost and Sander, 1993)) as the percentage of residues correctly predicted in three secondary structure states (helix, strand, other), the corresponding eight-state per-residue accuracy (<u>Q8</u>), the two-state per-protein accuracy (<u>Q2</u>) to describe the percentage of proteins correctly predicted as membrane-bound/water-soluble, and the ten-state per-protein accuracy (<u>Q10</u>) for the ten classes of localization. All numbers reported constituted averages over all proteins in the final test sets.

### Deep biology multi-task learning (DBMTL)

#### Concept

Thus far, successful applications of NLP techniques in computational biology (Alley, et al., 2019; Heinzinger, et al., 2019; Howard and Ruder, 2018; Rives, et al., 2019) used a two-step approach: Step 1: train a self-supervised model on protein sequences without annotations to learn vector representations (embeddings). Step 2: input embeddings to train supervised models on annotated data (e.g. secondary structure or localization). DBMTL merged the two steps into one single end-to-end model. This was achieved by applying a joint loss consisting of the four supervised tasks ((I) 3- and (II) 8-state secondary structure prediction, (III) 10-state per-protein localization and (IV) 2-state per-protein membrane/globular) and (V) the semi-supervised language modeling task. Thereby, the model could learn biochemical and biophysical properties of amino acids together (I, II & V) with the global properties of the macromolecules they constitute (III & IV). The idea was borrowed from the NLP transformer model (Vaswani, et al., 2017). These models consist of two building blocks, namely an encoder and a decoder. Here, we tested only the first approach (decoder only (Radford, et al., 2019)).

#### Architecture and training

The model used for DBMTL consists only of the decoder part of the transformer model (Fig. 1a: training, Fig. 1b: prediction/testing). We used 12 attention layers with maximum length of 1024 for inputs and outputs. The number of hidden layers was set to 1024, while the number of heads was set to 16. A dropout rate of 0.2 reduced the risk of over-fitting. For this model, the input differed between training and prediction (Fig. 1). During training, both the input and the output of each task were concatenated inserting a control token between them. During prediction, the model saw only the input followed by the control token. Each task had its own control token allowing the model to learn what to predict next: *<UNIREF50>* (Language Modelling), *<SSP3>* and <SSP8> (secondary structure prediction in 3 and 8 states), <LOC> (localization), <MS> (membrane-bound vs. water-soluble proteins).

**Fig. 1:**
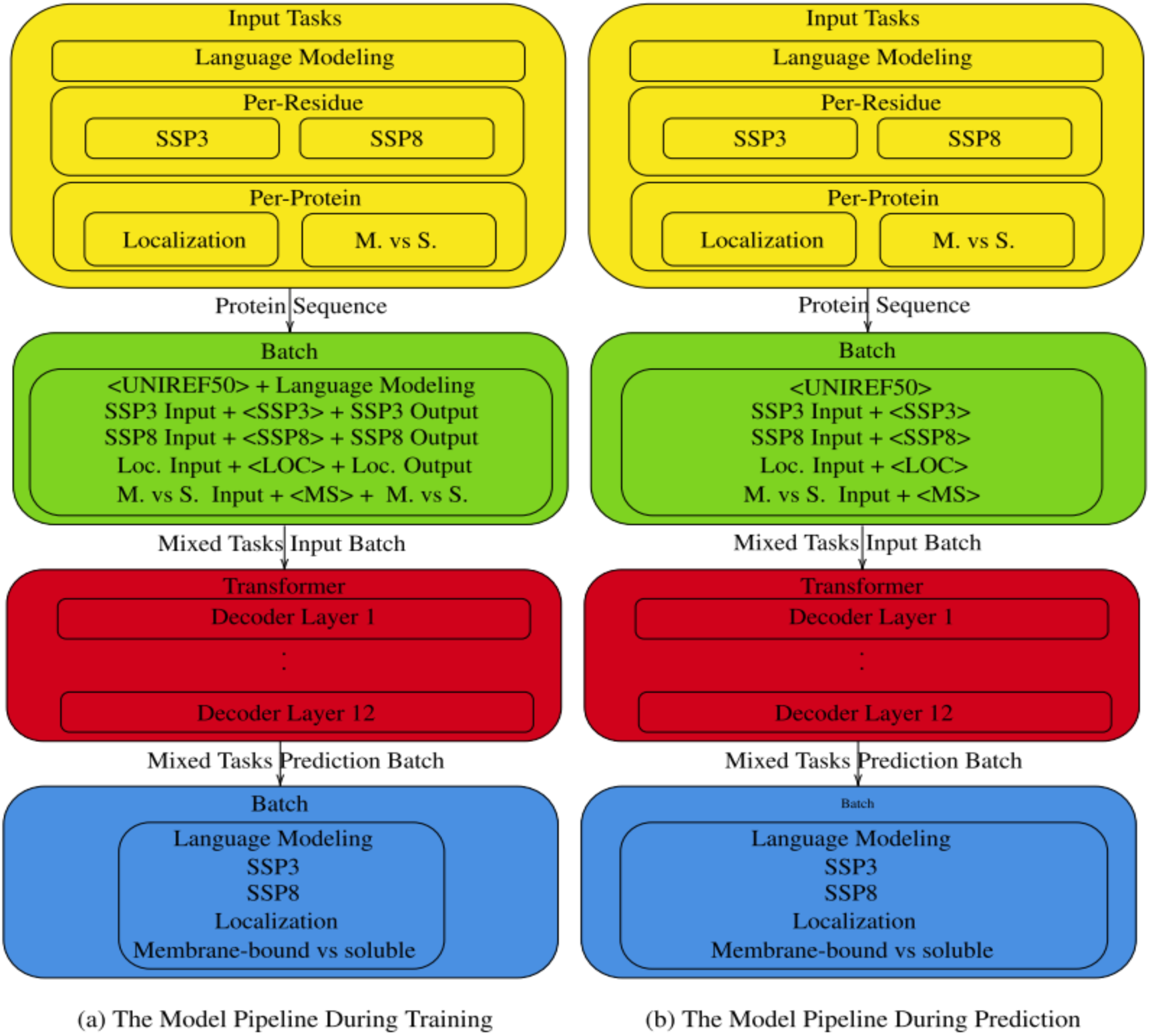
Sketch of DBTML architecture. (a) shows the model during training time and (b) shows the model during prediction time. The yellow block shows that the model supports the input from 5 different tasks. The green block is the only different block between training and prediction, during training it concatenate both input and output separated by a special control token, while during prediction it only append to the input the special control token. The red block shows the transformer model which contains 12 layers using only the decoder. The blue block shows that the output could be a mixed of different tasks output.

The model was trained on 6 Titan GPUs (local batch size: 512; global batch: 3072) using the Adam optimizer with 0.0002 learning rate, warm-up rate of 16k steps, and learning rate decay for 250k steps. The model was trained for a total number of 500K steps.

## Results

### Per-residue: secondary structure prediction competitive

DBMTL did not reach to the level of performance obtained by the top performers using evolutionary information from multiple sequence alignments (MSAs), such as NetSurfP-2.0 (Klausen, et al., 2019) or Spider3 (Heffernan, et al., 2017) and RaptorX (Wang, et al., 2016). However, it appeared to reach close and appeared to outperform other methods not using evolutionary information. In particular DeepSeqVec (Heinzinger, et al., 2019) and *ProtVec* (Asgari and Mofrad, 2015), the latter being one of the first approaches to successfully adopt NLP solutions (namely Google’s Word2vec (Mikolov, et al., 2013)) to problems in computational biology by extracting rules from languages that are context-independent. These results were not shown because we discovered a major problem with how Tensor2Tensor (T2T) (Vaswani, et al., 2018) utilizes the concept of teacher forcing for per-residue predictions: essentially, what works for the field of NLP or the per-protein prediction is not applicable. Instead, T2T automatically uses some of the observation for prediction (Discussion for detail). This problem did not affect the per-protein predictions.

### Per-protein performance high but not top

For both per-protein prediction tasks explored (localization and membrane-bound/water-soluble globular) DBMTL **<u>did not</u>** reach the top performance level (Fig. 2ab). For the prediction of localization, the best method DeepLoc reached Q10=78%, while the approach introduced here (DBMTL) remained nine percentage points below Q10=69% (Fig. 2a, Table SOM_3). DBMTL was numerically higher than other popular prediction methods using evolutionary information, namely (sorted by date): WoLF PSORT (Horton, et al., 2007), CELLO (Yu, et al., 2006), MultiLoc2 (Blum, et al., 2009), SherLoc2 (Briesemeister, et al., 2009), YLoc (Briesemeister, et al., 2010), iLoc-Euk (Chou, et al., 2011) and LocTree2 (Goldberg, et al., 2012). However, for localization prediction, the end-to-end solution DBMTL hardly outperformed the two-step approach realized earlier through SeqVec (Heinzinger, et al., 2019) embeddings (DeepSeqVec Fig. 2a). In contrast, the Word2vec-like approach ProtVec (Asgari and Mofrad, 2015) realized in DeepProtVec again performed much worse (over 20 percentage points: Fig. 2a).

**Fig. 2:**
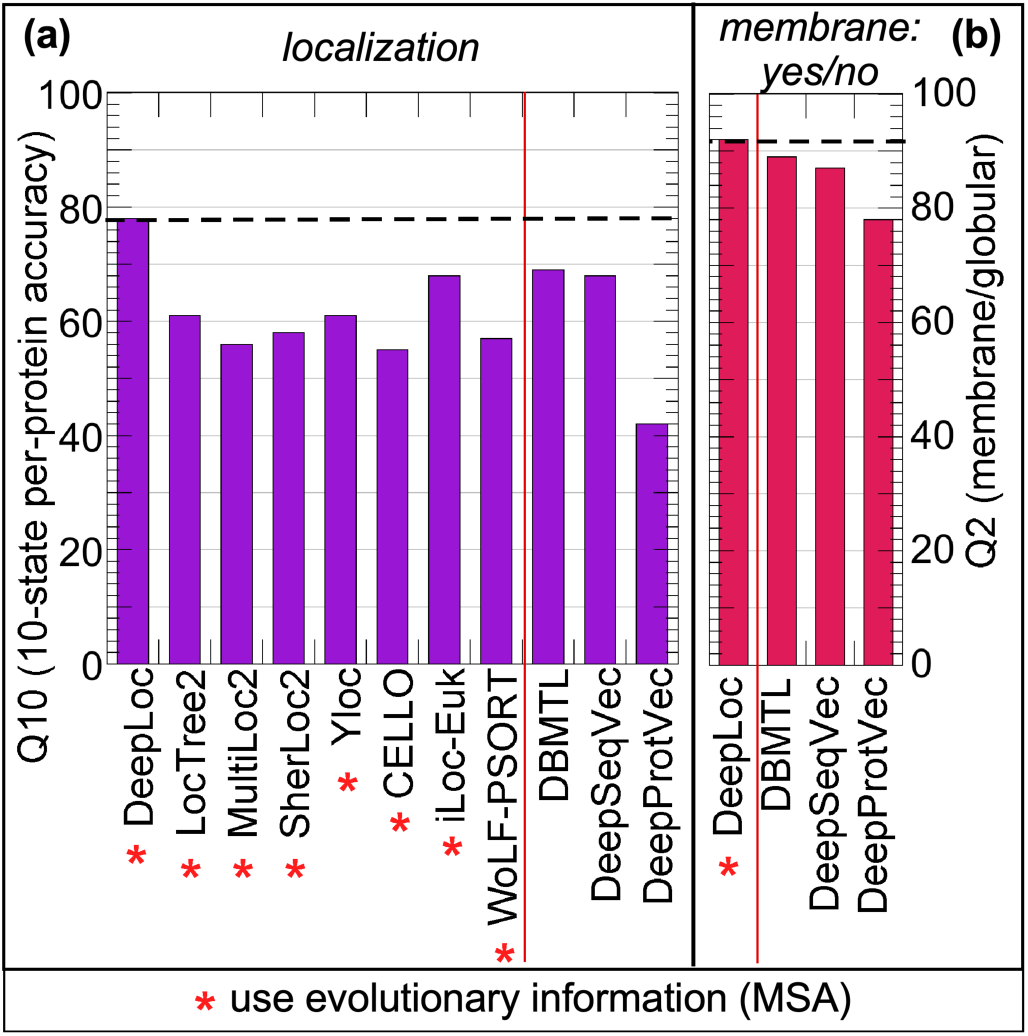
Performance comparison per-protein prediction. <u>Panel a:</u> The predictive power of DBMTL on the prediction of subcellular localization was also compared to top methods (numbers for Q10 taken from DeepLoc (Almagro Armenteros, et al., 2017) using evolutionary information. <u>Panel b:</u> The same data set as in Panel a was used to assess the predictive power of DBMTL for the classification of a protein into membrane-bound and water-soluble.

The situation was similar for the other per-protein task, namely the binary classification into membrane-bound and water-soluble, globular proteins. Although for this task, DBMTL missed the top level only by a smaller margin of three percentage points: Q2(DeepLoc)=92% vs. Q2(DBMTL)=89% (Fig. 2b, Table SOM_4). Once again, the novel end-to-end solution realized by DBMTL outperformed other solutions not using evolutionary information, e.g. the two-stage approach DeepSeqVec and the non-contextualized transfer-learning approach DeepProtVec by (Fig. 2b).

### Blazingly fast predictions

One of the main advantages of the transformer model over LSTM-based models such as ELMo (Peters, et al., 2018) is speed reached by processing input tokens in parallel through self-attention mechanisms (Bahdanau, et al., 2014). Using a single 1080Ti GPU, the model took ∼35 seconds to load, and with a batch size of 32, the prediction for a 1024-residue protein took, on average, 0.033 seconds. This was similar to the LSTM-based transfer-learning model SeqVec (Heinzinger, et al., 2019). In comparison to methods that use evolutionary information and have to first build the MSA, DBMTL reached a speed-up of approximately 110-fold compared to NetSurfP-2.0 using MMSeqs2 for creating alignments.

## Discussion

### Reporting negative results

Since the submission of the first version of this work, the authors spent all their resources on making the model openly available for the community to use. After trying to do so using machine learning toolkits (T2T (Vaswani, et al., 2018)), and failing to obtain speedy fixes by the community, the authors decided to re-engineer the underlying deep learning model. During the process of re-engineering the model, the authors discovered a fundamental problem with how the model calculates loss on secondary structure predictions, which undermines the authors’ confidence in the results initially reported. Since re-engineering the model with an open system and reproducing all experiments and results is time demanding and in-progress, the authors considered it important to update the initial version of the manuscript by removing results that are no longer sustained, until these can safely be verified or nullified. Additionally, the authors considered it important to update the preprint describing the shortfalls that emerged during re-engineering, so as to support fellow researchers not to commit the same mistakes.

### Flawed per-residue predictions

The core of DBMTL consists of the decoder part of a transformer model (Vaswani, et al., 2018; Vaswani, et al., 2017). Essentially, this model applies a stack of self-attention layers (Bahdanau, et al., 2014) followed by a linear transformation to predict the next token in a sequence, given all previous tokens in this sequence (self-supervision). These tokens do not necessarily need to come from the same language as for example in machine translation (Bahdanau, et al., 2014). Feeding a sentence concatenated with its translation to a self-supervised model like DBMTL or SeqVec (Heinzinger, et al., 2019) allows the translation to be conditioned upon the source language and in addition also upon those parts of the sentence that had already been translated. This produces more coherent translations and resembles the idea of teacher forcing (Williams and Zipser, 1989). Teacher forcing is used for problems which require sequential output generation like question answering or machine translation. During training, teacher forcing randomly uses predictions of previous states or ground truth labels which improves generalization. During inference, however, it relies solely on predictions as no translation exists, yet. The problem of the latter approach is efficiency: it requires LSTM-like sequential processing of a sentence where one token is predicted at a time in order to be able to give predictions of previous time steps to the model. With transformer models and their parallel processing capabilities being on the rise in the field of NLP, this time-consuming step was removed from the framework used here by using only ground truth labels for teacher forcing during training and evaluation/testing. Using this approach during evaluation approximates the actual performance during inference. However, this approximation might lead to overestimation because during inference, the model will have to rely on its own predictions instead of the high-quality ground truth data.

Applying the basic concept of translation and teacher forcing to protein sequences and their ‘translation’ to secondary structure, highlighted a pitfall of this tradeoff between reliability and computational overhead: the inherently well-structured nature of protein secondary structure allowed the model to reach state-of-the-art performance by always only replicating the secondary structure element of the previous residue. By feeding only ground truth labels, the model will just learn to replicate this annotation, since most secondary structure elements naturally occur in patches. Put simply, this is equal to shifting all ground truth labels in the secondary structure annotation by one, which in fact results in 85% 3-state secondary structure prediction performance. As a result, during inference the model will simply replicate the secondary structure prediction for the first residue because the model never learnt to switch from one secondary structure state to the other but just to replicate the teacher forcing signal. Obviously, the results are over-estimated performance values for all methods using the type of teacher forcing described here, namely all our implementations of per-residue predictions.

The problem might be addressed through at least two different approaches: remove teacher forcing at the cost of less coherent predictions, or use a model based on the encoder side of the transformer, e.g. Bert (Devlin, et al., 2018). The latter would replace the auto-regressive next-token prediction by an autoencoder-like training which tries to reconstruct corrupted or masked input.

### Limited gain for per-protein prediction tasks

For both per-protein tasks explored (localization and membrane-bound/not), the best competitors using evolutionary information still performed substantially better than the novel end-to-end solution (DBMTL; Fig. 2ab). For localization prediction, our approach still outperformed many popular solutions that use MSA (Fig. 2a). However, a recent publication might suggest this view to be possibly slightly distorted by the choice of data set and number of states (Savojardo, et al., 2018). Another reason for this result could be attributed to the technical limitation forcing proteins to be chopped to 1024 residues (using only the first and last 512 residues for protein longer than 512 amino acids). This clearly affected the per-protein data sets more than the per-residue data sets as the latter were taken from the PDB (Consortium, 2018), which tends to contain domain-like, aka. shorter fragments of long proteins. However, the same pre-processing (cutting long proteins) was performed for the classification into membrane-bound/not, whilst here performance was not as much below the top-performer (DeepLoc). This could be explained by two observations. Firstly, the signal of membrane-bound proteins is among the most dominant signals captured during language modeling (Heinzinger, et al., 2019). Secondly, subcellular localization relies heavily on certain short sequence motifs which might be removed when chopping the sequences (Almagro Armenteros, et al., 2017).

Another possible explanation as to why the end-to-end solution DBMTL improved less over the two-step approach for localization and membrane classification w.r.t. secondary structure prediction might be the smaller data sets for the former. All per-protein tasks had many orders of magnitude fewer samples than the per-residue tasks. Possibly, these were too few to tap into the full potential of the end-to-end solutions with many free parameters.

### How can semi-supervised multi-task learning be so successful?

Two factors may have contributed to the impressive improvement in secondary structure prediction: (1) semi-supervised learning and (2) model type. (1) *Semi-supervised learning*: previously, models tapping into the power of transfer learning capabilities of self-supervised language modeling might have pushed what can be achieved from single sequences to the limit without end-to-end training, i.e. coupling the pre-training phase and the task-specific fine-tuning phase. The DBMTL model tapped into the possibility of leveraging large amounts of unlabeled sequence (big data) while transferring knowledge between annotated proteins and proteins without any labels. This might have allowed the model to transfer the knowledge not only from the language model task to other tasks, but between tasks. (2) *Model type*: In NLP, the transformer model is currently achieving state-of-the-art results in various sequence to sequence problems (Dai, et al., 2019; Devlin, et al., 2018; Yang, et al., 2019). Leveraging this concept for biological sequences helped to push the boundaries of methods which do not rely on evolutionary information. The core module of all transformer models is the attention mechanism. This mechanism allows transformers to learn which regions in the input space (aka. sequence) contribute most to a prediction at a certain position (residue). Simply put, each output of the self-attention module is a weighted sum over all its inputs. One crucial advantage of this approach is that the length of the computational graph computing the dependency between any two residue positions i and j in a protein is independent of their sequential distance |i-j|, allowing to better capture long-range dependencies (Dai, et al., 2019).

### Speed-up

Most importantly, DBMTL speeds up over other state-of-the-art methods by not having to compile MSAs. How much time is gained through this step depends on how the MSAs are built. Compared to NetSurfP-2.0 we increase the speed ∼3000x (HHBlits3 is used to compute alignments) – 110x (MMseqs2 (Steinegger and Söding, 2017) is used to compute the alignments) On top, the advantage of using alignment-free predictors is growing hand-in-hand with the growth of bio-databases. These figures explain why, for many applications, even slightly less accurate, but faster predictors might be preferable to users when the top predictors become inexecutable due to limited computing resources.

## Conclusion

We presented a new method, DBMTL (Fig. 1), realizing end-to-end multi-task (MT) with the objective to push machine learning to a level at which it can compete with methods using evolutionary information in an alignment-free manner. DBMTL demonstrated how semi-supervised learning can be applied to the language of life, distilled in the form of protein sequences. By fusing self-supervised language modelling and supervised fine-tuning into one model which is trained with a joint loss that characterizes various protein properties, knowledge is transferred between different tasks, allowing the model to build a multi-modal understanding of proteins. In order to share as much knowledge between the tasks as possible, while still enabling the network to learn task-specific features, the MT implementation used in this work provides task-specific input signals. We implemented four supervised tasks: two on the level of residues (word-level), namely secondary structure prediction in 3- and 8-states, and two on the level of proteins (sentence-level), namely subcellular localization prediction and the classification membrane-vs-soluble proteins. For the time being, the performance of the per-residue (word-level) tasks cannot be trusted due to problems arising from adopting T2T (Vaswani, et al., 2018; Vaswani, et al., 2017) to this problem (Discussion). For the per-protein (sentence-level) tasks, DBMTL performed similar to many methods using evolutionary information from MSAs, however, it remained substantially below the level of today’s top performers (e.g. DeepLoc (Almagro Armenteros, et al., 2017)). Nevertheless, this is achieved without the costly step of building MSAs, thus our new method speeds up over competitors by about two orders of magnitude. Thus, semi-supervised MT approaches, as the one described in this work, may become a new frontier in computational biology and medicine, in particular when a wealth of unannotated data (sequences) contrasts with very constrained annotations (about protein structure and function).

## Abbreviations used

1D: one-dimensional – information representable in a string such as secondary structure or solvent accessibility;
3D: three-dimensional;
3D structure: three-dimensional coordinates of protein structure;
DBMTL: Deep Biology Multi-Task Learning;
NLP: Natural Language Processing;
PIDE: percentage of pairwise identical residues;

## Acknowledgements

The authors thank primarily to Tim Karl and Jian Kong for invaluable help with hardware and software and to Inga Weise for support with many other aspects of this work. We gratefully acknowledge the support of NVIDIA Corporation with the donation of two Titan GPUs used for this research. We also want to thank the LRZ (Leibniz Rechenzentrum) for providing us access to DGX-V1. This work was supported by a grant from the Alexander von Humboldt foundation through the German Ministry for Research and Education (BMBF: Bundesministerium fuer Bildung und Forschung) as well as by a grant from Deutsche Forschungsgemeinschaft (DFG– GZ: RO1320/4–1). Last, not least, thanks to all those who deposit their experimental data in public databases, and to those who maintain these databases.

## References

Abriata, L.A., et al. Assessment of hard target modelling in CASP12 reveals an emerging role of alignment-based contact prediction methods. Proteins: Structure, Function, and Bioinformatics 2018;86:97–112.

Alley, E.C., et al. Unified rational protein engineering with sequence-based deep representation learning. Nat Methods 2019:1–8.

Almagro Armenteros, J.J., et al. DeepLoc: prediction of protein subcellular localization using deep learning. Bioinformatics 2017;33(24):4049.

Asgari, E. and Mofrad, M.R. Continuous Distributed Representation of Biological Sequences for Deep Proteomics and Genomics. PloS one 2015;10(11):e0141287.

Bahdanau, D., Cho, K. and Bengio, Y. Neural machine translation by jointly learning to align and translate. arXiv preprint 1409.0473 2014.

Berman, H.M., et al. The protein data bank. Nucleic acids research 2000;28(1):235–242.

Blum, T., Briesemeister, S. and Kohlbacher, O. MultiLoc2: integrating phylogeny and Gene Ontology terms improves subcellular protein localization prediction. BMC Bioinformatics 2009;10:274.

Boutet, E., et al. UniProtKB/Swiss-Prot, the Manually Annotated Section of the UniProt KnowledgeBase: How to Use the Entry View. Methods Mol Biol 2016;1374:23–54.

Briesemeister, S., et al. SherLoc2: a high-accuracy hybrid method for predicting subcellular localization of proteins. J Proteome Res 2009;8(11):5363–5366.

Briesemeister, S., Rahnenfuhrer, J. and Kohlbacher, O. YLoc - an interpretable web server for predicting subcellular localization. Nucleic Acids Res 2010;38 Suppl:W497–502.

Capriotti, E., Fariselli, P. and Casadio, R. I-Mutant2.0: predicting stability changes upon mutation from the protein sequence or structure. Nucleic Acids Res 2005;33(Web Server issue):W306–310.

Casadio, R., Martelli, P.L. and Pierleoni, A. The prediction of protein subcellular localization from sequence: a shortcut to functional genome annotation. Brief Funct Genomic Proteomic 2008;7(1):63–73.

Chou, K.C., Wu, Z.C. and Xiao, X. iLoc-Euk: a multi-label classifier for predicting the subcellular localization of singleplex and multiplex eukaryotic proteins. PloS one 2011;6(3):e18258.

Consortium, U. UniProt: the universal protein knowledgebase. Nucleic acids research 2018;46(5):2699.

Cuff, J.A. and Barton, G.J. Evaluation and improvement of multiple sequence methods for protein secondary structure prediction. Proteins: Structure, Function, and Genetics 1999;34(4):508–519.

Dai, Z., et al. Transformer-xl: Attentive language models beyond a fixed-length context. arXiv preprint 1901.02860 2019.

Devlin, J., et al. Bert: Pre-training of deep bidirectional transformers for language understanding. arXiv preprint 1810.04805 2018.

Drozdetskiy, A., et al. JPred4: a protein secondary structure prediction server. Nucleic acids research 2015;43(W1):W389–W394.

Frishman, D. and Argos, P. Knowledge-based protein secondary structure assignment. Proteins: Structure, Function, and Genetics 1995;23:566–579.

Fu, L., et al. CD-HIT: accelerated for clustering the next-generation sequencing data. Bioinformatics 2012;28(23):3150–3152.

Goldberg, T., Hamp, T. and Rost, B. LocTree2 predicts localization for all domains of life. Bioinformatics 2012;28(18):i458–i465.

Goldberg, T., et al. LocTree3 prediction of localization. Nucleic Acids Res 2014;42(Web Server issue):W350–355.

Hamp, T. and Rost, B. Evolutionary profiles improve protein-protein interaction prediction from sequence. Bioinformatics 2015;31(12):1945–1950.

Hayat, S., et al. All-atom 3D structure prediction of transmembrane β-barrel proteins from sequences. Proceedings of the National Academy of Sciences 2015;112(17):5413–5418.

Heffernan, R., et al. Capturing non-local interactions by long short-term memory bidirectional recurrent neural networks for improving prediction of protein secondary structure, backbone angles, contact numbers and solvent accessibility. Bioinformatics 2017;33(18):2842–2849.

Heinzinger, M., et al. Modeling aspects of the language of life through transfer-learning protein sequences. BMC Bioinformatics 2019;20:723.

Heinzinger, M., et al. Modeling the Language of Life-Deep Learning Protein Sequences. bioRxiv 2019:614313.

Horton, P., et al. WoLF PSORT: protein localization predictor. Nucleic Acids Res 2007;35(Web Server issue):W585–587.

Howard, J. and Ruder, S. Universal language model fine-tuning for text classification. arXiv preprint 1801.06146 2018.

Jones, D.T. Protein secondary structure prediction based on position-specific scoring matrices. Journal of Molecular Biology 1999;292(2):195–202.

Kabsch, W. and Sander, C. Dictionary of protein secondary structure: pattern recognition of hydrogen bonded and geometrical features. Biopolymers 1983;22:2577–2637.

Klausen, M.S., et al. NetSurfP-2.0: Improved prediction of protein structural features by integrated deep learning. Proteins 2019.

Li, W. and Godzik, A. Cd-hit: a fast program for clustering and comparing large sets of protein or nucleotide sequences. Bioinformatics 2006;22(13):1658–1659.

Mehta, P.K., Heringa, J. and Argos, P. A simple and fast approach to prediction of protein secondary structure from multiply aligned sequences with accuracy above 70%. Protein Science 1995;4:2517–2525.

Mikolov, T., et al. Efficient estimation of word representations in vector space. ArXiv 2013:1301.3781.

Nair, R. and Rost, B. Better prediction of sub-cellular localization by combining evolutionary and structural information. Proteins: Structure, Function, and Bioinformatics 2003;53(4):917–930.

Perdigao, N., et al. Unexpected features of the dark proteome. Proceedings of the National Academy of Sciences of the United States of America 2015.

Peters, M.E., et al. Deep contextualized word representations. arXiv 2018;1802.05365.

Pollastri, G., et al. Prediction of coordination number and relative solvent accessibility in proteins. Proteins 2002;47(2):142–153.

Radford, A., et al. Language models are unsupervised multitask learners. OpenAI Blog 2019;1(8).

Radivojac, P., et al. Protein flexibility and intrinsic disorder. Protein Science 2004;13:71–80.

Rives, A., et al. Biological structure and function emerge from scaling unsupervised learning to 250 million protein sequences. bioRxiv 2019:622803.

Rost, B. Protein secondary structure prediction continues to rise. Journal of Structural Biology 2001;134:204–218.

Rost, B., et al. Transmembrane helix prediction at 95% accuracy. Protein Science 1995;4:521–533.

Rost, B. and Sander, C. Improved prediction of protein secondary structure by use of sequence profiles and neural networks. Proceedings of the National Academy of Sciences 1993;90:7558–7562.

Rost, B. and Sander, C. Prediction of protein secondary structure at better than 70% accuracy. Journal of Molecular Biology 1993;232:584–599.

Rost, B. and Sander, C. Combining evolutionary information and neural networks to predict protein secondary structure. Proteins: Structure, Function, and Genetics 1994;19:55–72.

Rost, B. and Sander, C. Conservation and prediction of solvent accessibility in protein families. Proteins: Structure, Function, and Genetics 1994;20(3):216–226.

Savojardo, C., et al. BUSCA: an integrative web server to predict subcellular localization of proteins. Nucleic Acids Res 2018;46(W1):W459–W466.

Steinegger, M., et al. HH-suite3 for fast remote homology detection and deep protein annotation. BMC Bioinformatics 2019;20(1):473.

Steinegger, M., Mirdita, M. and Söding, J. Protein-level assembly increases protein sequence recovery from metagenomic samples manyfold. Nat Methods 2019:1.

Steinegger, M. and Söding, J. MMseqs2 enables sensitive protein sequence searching for the analysis of massive data sets. Nature biotechnology 2017;35(11):1026.

Suzek, B.E., et al. UniRef clusters: a comprehensive and scalable alternative for improving sequence similarity searches. Bioinformatics 2015;31(6):926–932.

Vaswani, A., et al. Tensor2Tensor for neural machine translation. arXiv 2018;1803.07416.

Vaswani, A., et al. Attention is all you need. In, Advances in neural information processing systems. 2017. p. 5998–6008.

Velankar, S., et al. SIFTS: structure integration with function, taxonomy and sequences resource. Nucleic acids research 2012;41(D1):D483–D489.

Wang, G. and Dunbrack Jr, R.L. PISCES: a protein sequence culling server. Bioinformatics 2003;19(12):1589–1591.

Wang, S., et al. RaptorX-Property: a web server for protein structure property prediction. Nucleic acids research 2016;44(W1):W430–W435.

Williams, R. and Zipser, D. A learning algorithm for continually running fully recurrent neural networks. Neural Computation 1989;1:270–280.

Yang, Y., et al. Sixty-five years of the long march in protein secondary structure prediction: the final stretch? Briefings in bioinformatics 2016;19(3):482–494.

Yang, Z., et al. XLNet: Generalized Autoregressive Pretraining for Language Understanding. arXiv preprint 1906.08237 2019.

Yu, C.S., et al. Prediction of protein subcellular localization. Proteins 2006;64(3):643–651.

Zhang, Q.C., et al. Structure-based prediction of protein-protein interactions on a genome-wide scale. Nature 2012;490(7421):556–560.

